# Domain-selective BET inhibition attenuates transcriptional and behavioral responses to cocaine

**DOI:** 10.1101/2021.10.04.463083

**Authors:** MB Singh, CJ Babigian, GC Sartor

## Abstract

Epigenetic pharmacotherapies have emerged as a promising treatment option for substance use disorder (SUD) due to their ability to reverse maladaptive transcriptional and behavioral responses to drugs of abuse. In particular, inhibitors of bromodomain and extra terminal domain (BET) reader proteins have been shown to reduce cocaine- and opioid-seeking behaviors in rodents. However, only pan-BET inhibitors, small molecules that bind to both bromodomains (BD1 and BD2) with all BET proteins, have been investigated in animal models of SUD. Given the potential side effects associated with pan-BET inhibitors, safer and more selective strategies are needed to advance BET therapeutics as a potential treatment for SUD. Here, we show that RVX-208, a clinically tested, BD2-selective BET inhibitor, dose-dependently reduced cocaine conditioned place preference in male mice, similar to the pan-BET inhibitor JQ1. In other behavioral experiments, RVX-208 treatment did not alter distance traveled, anxiety-like behavior, or novel object recognition memory. At the transcriptional level, RVX-208 attenuated the expression of multiple cocaine-induced genes in the nucleus accumbens. RVX-208 produced a distinct transcriptional response in stimulated primary neurons compared to JQ1 but had little effect on gene expression in non-stimulated neurons. Together, these data indicate that targeting domain-specific BET mechanisms may be an effective and safer strategy to reduce cocaine-induced neurobehavioral adaptations.

## Introduction

Repeated drug use alters histone acetylation processes in reward-related brain regions, which contributes, in part, to the molecular maladaptations involved in drug intake and relapse (Rogge and Wood, 2013). While writers (e.g., histone acetyltransferases, HATs) and erasers (e.g., histone deacetylases, HDACs) of histone acetylation have well-established roles in animal models of substance use disorder (SUD) (Kennedy et al., 2013; Malvaez et al., 2013, 2011; Rogge et al., 2013), readers of histone acetylation, called bromodomains, have also emerged as potential therapeutic targets (Egervari et al., 2017; Guo et al., 2020; Sartor et al., 2015). Specifically, pharmacological inhibition of bromodomain and extra terminal domain (BET) reader proteins (BRD2, BRD3, BRD4 and BRDT) has been shown to normalize transcriptional aberrations and behavioral symptoms in a wide range of disease models, including SUD (For review see: Singh and Sartor, 2020; Zaware and Zhou, 2017). Recognizing the growing success and usefulness of BET inhibitors, many large and small pharmaceutical companies have developed new and more selective BET therapeutics that are being tested in a variety of preclinical and clinical studies (Lu et al., 2020).

Each BET protein contains highly conserved, tandem bromodomains (BD1 and BD2) that have been successfully targeted by small molecule inhibitors (Wu et al., 2019). First-generation BET inhibitors, also referred to as pan-BET inhibitors (e.g., JQ1, IBET151, IBET762), displayed similar affinity for BD1 and BD2 within all BET proteins and have been shown to alter disease-associated gene expression in multiple cell types by blocking BET bromodomain interactions with acetylated histones (Dawson et al., 2011; Filippakopoulos et al., 2010; Huang et al., 2009; Nicodeme et al., 2010). In preclinical SUD models, the pan-BET inhibitor, JQ1, was found to attenuate behavioral and transcriptional responses to cocaine and heroin in mice and rats (Egervari et al., 2017; Guo et al., 2020; Sartor et al., 2015). However, our understanding of bromodomain-specific BET mechanisms in drug-seeking behaviors remains limited, as JQ1 blocks both bromodomains (BD1 and BD2) within all BET proteins. Additionally, pan-BET inhibitors similar to JQ1 are known to have side effects in humans that may limit their clinical applications for SUD (Amorim et al., 2016; Berthon et al., 2016).

To reduce potential toxic liabilities associated with pan-BET inhibitors, more recent efforts have focused on developing domain-selective BET inhibitors that target specific bromodomains within BET proteins (Petretich et al., 2020). RVX-208 (also called Apabetalone and RVX000222) was one of the first small molecule BET inhibitors to display preferential binding to the BD2 domain within BET proteins (Picaud et al., 2013). In cell lines, RVX-208 only modestly affected BET-dependent gene expression and produced a unique transcriptional outcome compared to JQ1 (Picaud et al., 2013). More recent studies indicate that RVX-208 may offer therapeutic benefits in multiple disease models by blocking the transcriptional responses to inflammatory stimuli (Tsujikawa et al., 2019; Wasiak et al., 2020). In humans, RVX-208 was found to be safe and well tolerated, and it is currently being investigated in Phase II/III clinical trials for the treatment of acute coronary syndromes, atherosclerosis, and Alzheimer’s disease (McNeill, 2010; Nikolic et al., 2015). Given that RVX-208 is a clinically tested, potentially safe, BD2-selective BET inhibitor, we sought to investigate its effects on cocaine-induced behavior and transcriptional activity. Here, we found that RVX-208 dose-dependently reduced cocaine conditioned place preference without altering locomotor activity, thigmotaxis, and novel object recognition learning. At the transcriptional level, RVX-208 reduced cocaine-induced gene expression in the nucleus accumbens (NAc) and produced a disparate transcriptional response in primary neurons compared to JQ1. Together, these data indicate that the BET BD2 domain plays an important role in the behavioral and transcriptional responses to cocaine and that BD2 BET inhibition has limited effects in non-stimulated neurons and other behaviors.

## Materials and Methods

### Drug treatments

For *in vitro* studies, JQ1 and RVX-208 (Cayman chemicals) were dissolved in dimethyl sulfoxide (DMSO) and administered at 0.1% v/v. In *in vivo* studies, JQ1 and RVX-208 were dissolved in 10% DMSO and 5% Tween 80 (v/v) and then diluted with sterile saline. RVX-208, JQ1, and vehicle were administered by an intraperitoneal (i.p.) injection at a volume of 0.1-0.15 mL. Cocaine HCl (10 mg/kg, National Institutes of Health’s National Institute on Drug Abuse Drug Supply Program) was dissolved in 0.9% sterile saline and administered by an i.p. injection.

### Animals

Male C57BL/6 mice (10-12 weeks old; Charles River Laboratories) were group housed under a reverse 12 h/12 h light/dark cycle and had *ad libitum* access to food and water. For primary neuron experiments, a pregnant female rat (Sprague Dawley; Charles River Laboratories) carrying embryonic day 16 (E16) pups was single housed with ad libitum access to food and water. Animals were housed in a humidity- and temperature-controlled and AAALAC-accredited animal facility at the University of Connecticut. Experiments were approved by the Institutional Animal Care and Use Committee and conducted according to guidelines established by the U.S. Public Health Service Policy on Humane Care and Use of Laboratory Animals.

### Primary neurons

Primary rat cortical neurons were isolated at E18 in cold 0.1 M PBS with 0.1% glucose wt./vol. Tissue was digested with papain for 10 min at 37ºC (2 mg/ml papain solution in Hibernate-E minus CaCl2, BrainBits LLC). Papain solution was removed, and the tissue was washed three times with Hibernate-E solution followed by 15-minute incubation in Hibernate-E solution and DNAse (1% wt/vol.) at room temperature. Cells were mechanically dissociated with a fire-polished pasteur pipette, resuspended in NbActive4 media (BrainBits LLC), and plated at a density of ∼400,000 cells on a poly-L-ornithine (100 μg/ml) and laminin (15 μg/ml) coated 12-well plates. Neurons were maintained at 5% CO_2_ and 37º C for 12 days in vitro (DIV), and half the media was replaced with fresh media every three days. In some experiments, neurons were treated with DMSO (vehicle), RVX-208 (100 nM and 500 nM) and JQ1 (100 nM and 500 nM) for 1 h, 2 h and 3 h. In other experiments, neurons received a 2 h cotreatment of AMPA (20 μM) or BDNF (30 ng/mL) with DMSO, RVX-208 (100 or 500 nM), or JQ1 (100 or 500 nM). These time points were selected to capture the early effects of BET inhibition and are similar to previous studies (Sartor et al., 2015; Sullivan et al., 2015). BDNF and AMPA stimulation were used because previous reports indicate that BET protein activity is regulated by glutamate and tyrosine receptor kinase B (TrKB) receptor signaling (Korb et al., 2015). After treatment, neurons were collected for RNA extraction as described below.

### RT-qPCR

In *in vivo* studies, mice received an i.p. injection of vehicle or RVX-208 (50 mg/kg) 5 minutes before an i.p. injection of saline or cocaine (10 mg/kg). The nucleus accumbens (NAc) was collected and homogenized in TRIzol 2 hours after the last injection. In cell culture experiments, the media was aspirated and 1 ml of TRIzol was added to each well. RNEasy Mini Kit (Qiagen) was used for RNA extraction according to the manufacturer’s instructions. RNA was reverse transcribed using High-Capacity cDNA Reverse Transcription Kit (Thermo Scientific). Using validated TaqMan primer probes for *Bdnf* (Thermo Scientific, Rn02531967_s1, Mm04230607_s1), *Gria1* (Rn06323759_m1, Mm00433752_m1), *Arc* (Rn00571208_g1, Mm01204954_g1), *Nr4a1* (Rn01533237_m1, Mm01300401_m1), *c-Fos* (Rn02396759_m1, Mm00487425_m1), and *Fosb* (Mm00500401_m1), cDNA was run in triplicate and analyzed using the 2−ΔΔCT method with β*-actin* (Rn00667869_m1, Mm02619580_g1) as a normalization control. The genes selected for measurement are known BET target genes and/or genes involved in cocaine-induced plasticity (Korb et al., 2015; Sartor et al., 2015; Sullivan et al., 2015). *Fosb* levels were extremely low in non-stimulated primary neurons (RT values > 35), and for this reason these data could not reported. The housekeeping gene selected for normalization was unchanged between treatments.

### Conditioned place preference (CPP)

The CPP apparatus (Med Associates Inc.) consisted of two compartments with distinct contextual cues on each side: black walls and grid rod-style floors in one compartment and white walls mesh style floors in the other compartment. The compartments were separated by a divider with a small door that could be opened or closed. In a pretest, drug-free acclimation session, mice were allowed to freely explore both compartments for 15 min via an opening in the partition. The time spent on each side of the CPP chamber was automatically recorded by infrared photobeam detectors on the chamber walls. Mice that exhibited a strong bias for one side of the chamber during the pretest (> 65% of time spent on one side) were excluded from further testing. For the next three consecutive days, animals were conditioned as previously described (Sartor et al., 2015; Sartor and Aston-Jones, 2012). During the 30-minute conditioning sessions, one side of the chamber was paired with an i.p. saline injection and the other side was paired with a cocaine injection (10 mg/kg, i.p.). Conditioning occurred in morning and afternoon sessions in a counterbalanced design (at least 4 h apart). Five minutes before each cocaine conditioning session, mice received an i.p. injection of vehicle, JQ1 (50 mg/kg) or RVX-208 (10, 25, or 50 mg/kg). One day after the last conditioning session, mice received a drug-free CPP test where they had free access to each side of the chamber. The preference score was calculated by subtracting the time spent on the cocaine-paired side during the posttest from the time spent on the same side during the pretest.

### Open field

The open field (Ugo Basile) consists of a 46 × 46 cm chamber with opaque gray walls. Baseline behavior (distance traveled and time spent in the inner and outer zone) was measured in a 30-minute, drug-free habituated test. Treatments were assigned so that baseline measurements did not differ between groups. The next day, mice were injected with vehicle or RVX-208 (50 mg/kg, i.p.) and were placed in the chamber 5 minutes post-injection. Distance traveled and time spent in the inner and outer zone were measured for 30 minutes using EthoVision tracking software (Noldus).

### Novel object recognition (NOR)

On days 1-3, mice were allowed to freely explore a clean mouse home cage (without objects) for 5 min per day. On day 4, mice were exposed to two identical objects in the cage for 10 min. After the test, mice were immediately injected with vehicle or RVX-208 (50 mg/kg) and returned to the colony room. Previous studies using this approach showed that JQ1 reduced NOR (Korb et al., 2015). The next day, mice were tested in a 10 min retention trial with one familiar object and one novel object. The objects consisted of a stack of Lego Duplo bricks and a 100 ml glass beaker. The location of the objects was counterbalanced for left and right positions in each treatment group. The objects were thoroughly cleaned with 70% ethanol after each session. Total time spent exploring and facing each object within 2.5 cm was calculated by Ethovision tracking software. Results were presented as time exploring the objects and discrimination index: [(novel object investigation time minus familiar object investigation time)/(total investigation time of both objects) * 100].

### Data analysis

Statistical analyses were performed with GraphPad Prism 7.0 software. Mean values from behavioral studies and Rq values from qRT-PCR experiments were compared between groups using analysis of variance (ANOVA) or Student’s t-test. When comparing multiple groups, a Bonferroni correction test was utilized. Data are expressed as means ±SEM, and the level of significance was set at *P* < 0.05.

## Results

### The effects of RVX-208 on gene expression in primary neurons

In *in vitro* experiments, primary neurons were treated with vehicle, RVX-208 (100 nM or 500 nM), or JQ1 (100 nM or 500 nM) and gene expression was measured after 1, 2, and 3 h of treatment (**Figures 1A-O**). RVX-208 had minimal effect on gene expression, with the only significant change occurring in *Bdnf* expression after 2 h of treatment (One-way ANOVA main effect for 2 h: F_(4,13)_ = 9.313, P = 0.0009) (**Figure 1B**). JQ1 on the other hand, altered the expression of most genes examined, including: a decrease in *Bdnf* at 1, 2 and 3 h timepoints (One-way ANOVA main effect for 1 h, F_(4,13)_ = 4.326, P = 0.0192; for 2 h, F_(4,13)_ = 9.313, P = 0.0009; for 3 h, F_(4, 13)_ = 6.881, P = 0.0033) (**Figures 1A-C**); a decrease in *Arc* at 1, 2 and 3 h timepoints (One-way ANOVA main effect for 1 h, F_(4,13)_ = 4.96, P = 0.012; for 2 h F_(4,13)_ = 5.13, P = 0.01; for 3 h, F_(4, 13)_ = 23.34, P < 0.0001) (**Figures 1D-F**); a decrease in *Nr4a1* at 1, 2 and 3 h timepoints (One-way ANOVA main effect for 1 h, F_(4,13)_ = 7.588, P = 0.0022; for 2 h, F_(4,13)_ = 14.23, P = 0.0001; for 3 h, F_(4, 13)_ = 27.55, P < 0.0001) (**Figures 1G-I**); and an increase in *c-Fos* at the 3 h timepoint (One-way ANOVA main effect for 3 h, F_(4, 13)_ = 10.55, P = 0.0005) (**Figure 1L**). RVX-208 and JQ1 did not affect *Gira1* expression (F values < 2.1, P values > 0.05) (**Figures 1M-O**).

**Figure 1:**
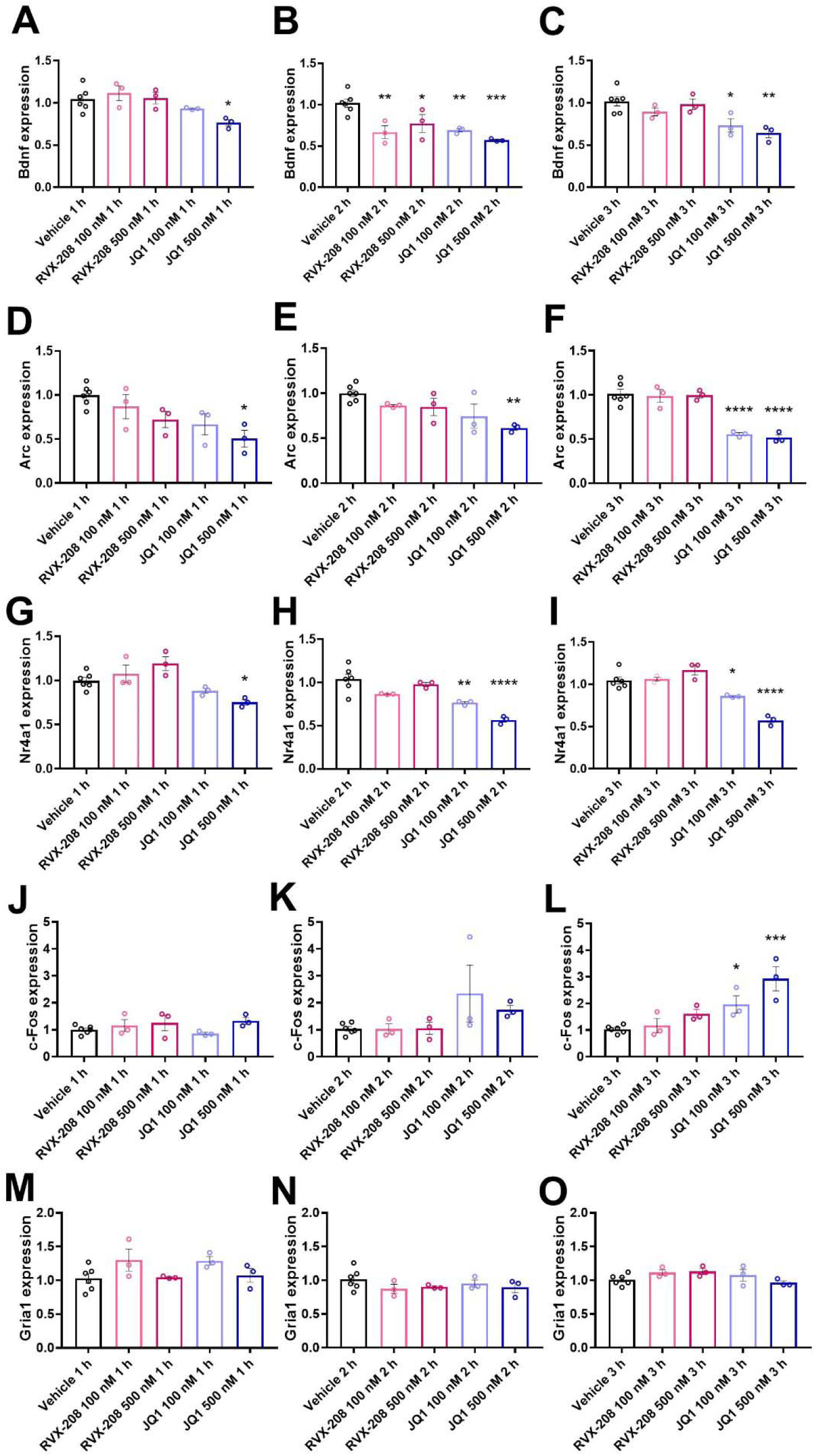
RVX-208 has limited effects in non-stimulated primary neurons. Primary neurons were treated with vehicle, RVX-208, or JQ1 (100 nM and 500 nM) for 1, 2, and 3 hours. Following treatment, *Bdnf* (A-C), *Arc* (D-F), *Nr4a1* (G-I), *c-Fos* (J-L), and *Gria1* (M-O) expression levels were measured via RT-qPCR. \**P* < 0.05, \*\**P* < 0.01, \*\*\**P* < 0.001, and \*\*\*\**P* < 0.0001 indicate a significant difference using Bonferroni *post hoc* test. Data are mean ±SEM.

### The effects of RVX-208 on gene expression in stimulated primary neurons

Previous experiments indicate that the BET BD2 domain is responsible for stimulus-induced gene expression in cell lines (Gilan et al., 2020). To examine if RVX-208 alters gene expression in stimulated neurons, we co-treated neurons for 2 h with vehicle, RVX-208 (500 nM), or JQ1 (500 nM) plus BDNF (**Figure 2**) or AMPA (**Figure 3**). In BDNF and AMPA stimulated neurons, all genes examined, except *Gria1*, were elevated by more than 5-fold compared to untreated neurons (baseline) (**Figures 2 and 3**). In BDNF simulated neurons, JQ1 but not RVX-208 significantly reduced the expression of *Bdnf* (One-way ANOVA main effect: F_(3,8)_ = 69.93, P < 0.0001 followed by Bonferroni post-hoc test, P < 0.0001 for BDNF vs. JQ1 + BDNF, P < 0.0001 for Vehicle + BDNF vs. JQ1 + BDNF) (**Figure 2A**). *Arc* expression was reduced by RVX-208, while JQ1 elevated *Arc* expression in BDNF stimulated neurons (One-way ANOVA main effect: F_(3,8)_ = 22.02, P = 0.0003 followed by Bonferroni post-hoc test, P = 0.0059 for BDNF vs. JQ1 + BDNF, P = 0.004 for Vehicle + BDNF vs. RVX-208 + BDNF) (**Figure 2B**). *Nr4a1* was reduced by RVX-208 but unchanged by JQ1 (One-way ANOVA main effect: F_(3,8)_ = 13.43, P = 0.0017 followed by Bonferroni post-hoc test, P = 0.0016 for BDNF vs. RVX-208 + BDNF, P = 0.0081 for vehicle + BDNF vs. RVX-208 + BDNF) (**Figure 2C**), and *c-fos* was significantly elevated by JQ1 but unchanged RVX-208 in BDNF stimulated neurons (One-way ANOVA main effect: F_(3,8)_ = 78.32, P < 0.0001 followed by Bonferroni post-hoc test, P < 0.0001 for BDNF vs. JQ1 + BDNF, P < 0.0001 for vehicle + BDNF vs. JQ1 + BDNF) (**Figure 2D**). Although Gria1 was not altered by BDNF stimulation compared to baseline, RVX-208 and JQ1 significant reduced *Gria1* expression in BDNF stimulated neurons (One-way ANOVA main effect: F_(3,8)_ = 14.08, P = 0.0015 followed by Bonferroni post-hoc test, P = 0.0148 for BDNF vs. JQ1 + BDNF, P = 0.0123 for Vehicle + BDNF vs. JQ1 + BDNF, P = 0.007 for BDNF vs. RVX-208 + BDNF, P = 0.006 for Vehicle + BDNF vs. RVX-208 + BDNF) (**Figure 2E**).

**Figure 2:**
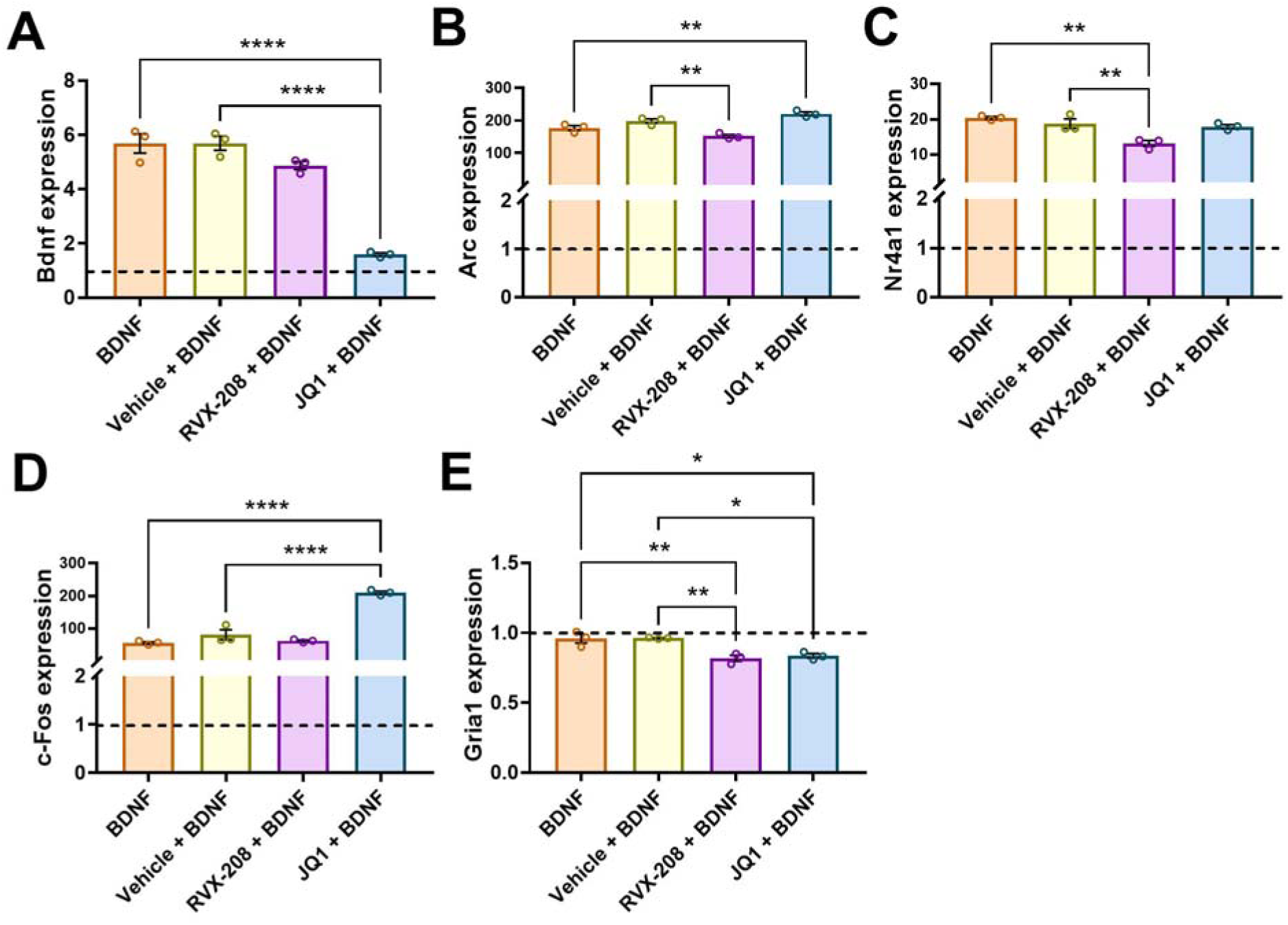
RVX-208 alters the expression of multiple genes in BDNF stimulated primary neurons. Primary neurons were co-treated with BDNF, vehicle + BDNF, RVX-208 + BDNF or JQ1 + BDNF for 2 hours. The expression of (A) *Bdnf*, (B) *Arc*, (C) *Nr4a1*, (D) *c-Fos*, and (E) *Gria1* was measured via RT-qPCR. Horizontal dotted line indicates baseline expression of untreated neurons. \**P* < 0.05, \*\**P* < 0.01, and \*\*\*\**P* < 0.0001 indicate a significant difference using Bonferroni *post hoc* test. Data are mean ±SEM.

**Figure 3:**
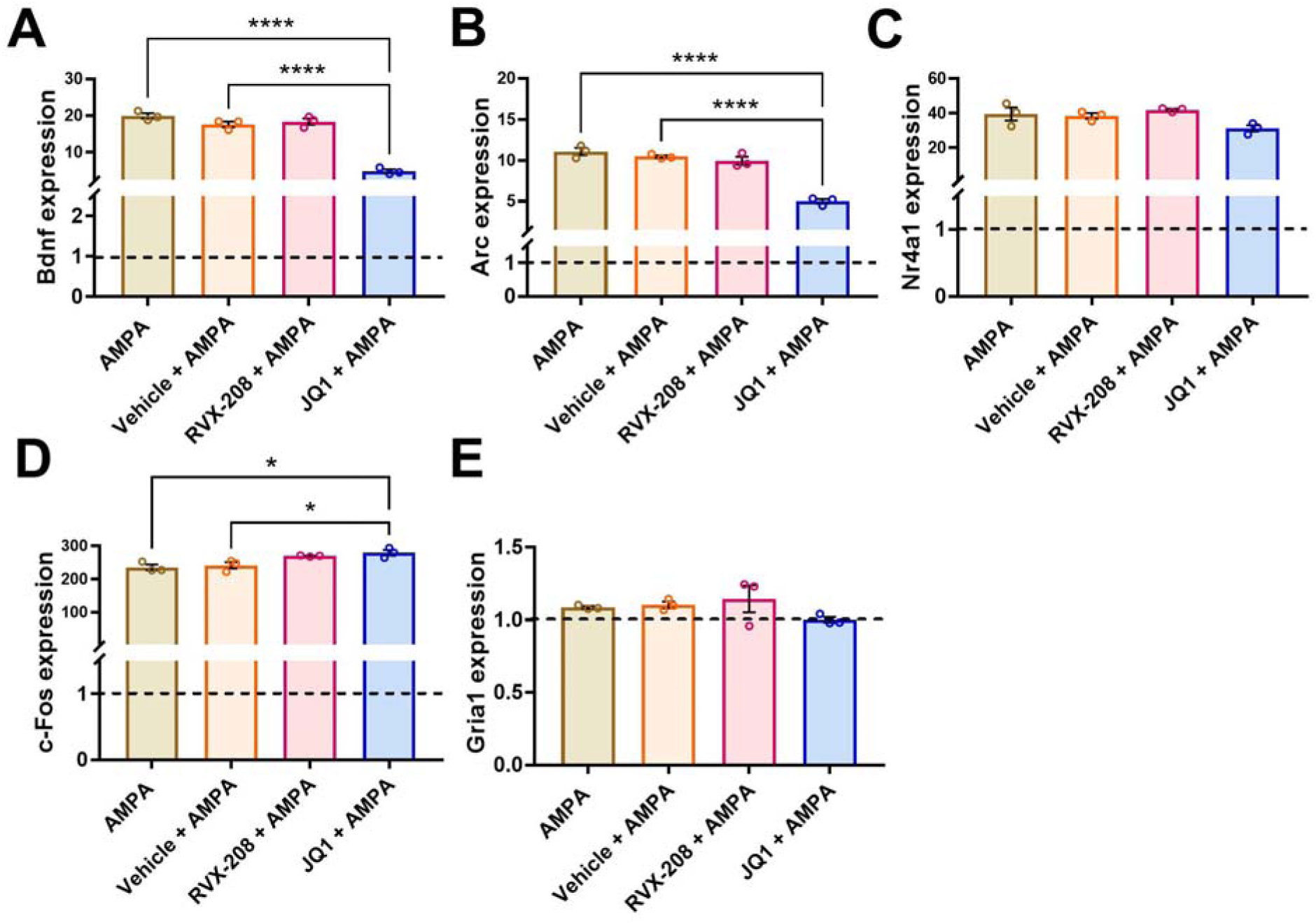
RVX-208 has no effect on gene expression in AMPA stimulated primary neurons. Primary neurons were co-treated with AMPA, vehicle + AMPA, RVX-208 + AMPA, or JQ1 + AMPA for 2 hours. The expression of (A) *Bdnf*, (B) *Arc*, (C) *Nr4a1*, (D) *c-Fos*, and (E) *Gria1* was measured via RT-qPCR. Horizontal dotted line indicates baseline expression of untreated neurons. \**P* < 0.05 and \*\*\*\**P* < 0.0001 indicate a significant difference using Bonferroni *post hoc* test. Data are mean ±SEM.

In AMPA stimulated neurons, RVX-208 did not affect the expression of the genes measured (P values > 0.05) (**Figure 3**). JQ1, on the other hand, decreased *Bdnf* (One-way ANOVA main effect: F_(3,8)_ = 90.82, P < 0.0001 followed by Bonferroni post-hoc test, P < 0.0001 for AMPA vs. JQ1 + AMPA, P < 0.0001 for vehicle + AMPA vs. JQ1 + AMPA) (**Figure 3A**), *Arc* (One-way ANOVA main effect: F_(3,8)_ = 58.86, P < 0.0001 followed by Bonferroni post-hoc test, P < 0.0001 for AMPA vs. JQ1 + AMPA, P < 0.0001 for vehicle + AMPA vs. JQ1 + AMPA) (**Figure 3B**), and increased *c-fos* expression under AMPA stimulation (One-way ANOVA main effect: F_(3,8)_ = 7.464, P = 0.0105 followed by Bonferroni post-hoc test, P = 0.0208 for AMPA vs. JQ1 + AMPA, P = 0.044 for vehicle + AMPA vs. JQ1 + AMPA) (**Figure 3D**), but had no effect on *Gria1* and *Nr4a1* expression (F values < 1.6, P values > 0.05) (**Figures 3C and E**).

### Effects of RVX-208 on cocaine-induced gene expression in the NAc

To examine the effects of RVX-208 on cocaine-induced gene expression, mice were treated with vehicle or RVX-208 (50 mg/kg) prior to an injection of cocaine (10 mg/kg) or saline (**Figure 4**). In the NAc, cocaine significantly increased the expression of *Arc* (One-way ANOVA main effect: F _(3, 28)_ = 11.39 P < 0.0001 followed by Bonferroni post-hoc test, P = 0.0005 for vehicle/saline vs. vehicle/cocaine) (**Figure 4B**), *Nr4a1* (One-way ANOVA main effect: F _(3, 28)_ = 9.171, P = 0.0002 followed by Bonferroni post-hoc test, P = 0.0002 for vehicle/saline vs. vehicle/cocaine) (**Figure 4C)**, *c-Fos* (One-way ANOVA main effect: F _(3, 28)_ = 12.12, P < 0.0001 followed by Bonferroni post-hoc test, P = 0.0002 for vehicle/saline vs. vehicle/cocaine) (**Figure 4D**), and *Fosb* (One-way ANOVA main effect: F _(3, 28)_ = 18.78, P < 0.0001 followed by Bonferroni post-hoc test, P < 0.0001 for vehicle/saline vs. vehicle/cocaine) (**Figure 4E**), but not *Gria1* (One-way ANOVA main effect: F _(3, 28)_ = 2.727, P = 0.0629) (**Figure 4F**). Although a statistically significant main effect was observed in *Bdnf* expression (One-way ANOVA main effect: F _(3, 28)_ = 3.142, P = 0.0409), no significant changes were observed in the post hoc comparisons (P values > 0.05) due to the high variability within and between each group (**Figure 4A**). When comparing vehicle/cocaine to RVX-208/cocaine, RVX-208 attenuated cocaine-induced gene expression of *Arc* (P = 0.0003), *Nr4a1* (P = 0.0125), *c-Fos* (P = 0.0002), and *Fosb* (P < 0.0001) (**Figures 4B-E**). There were no significant changes in gene expression when comparing vehicle/saline to RVX-208/saline treated mice (P values > 0.05).

**Figure 4:**
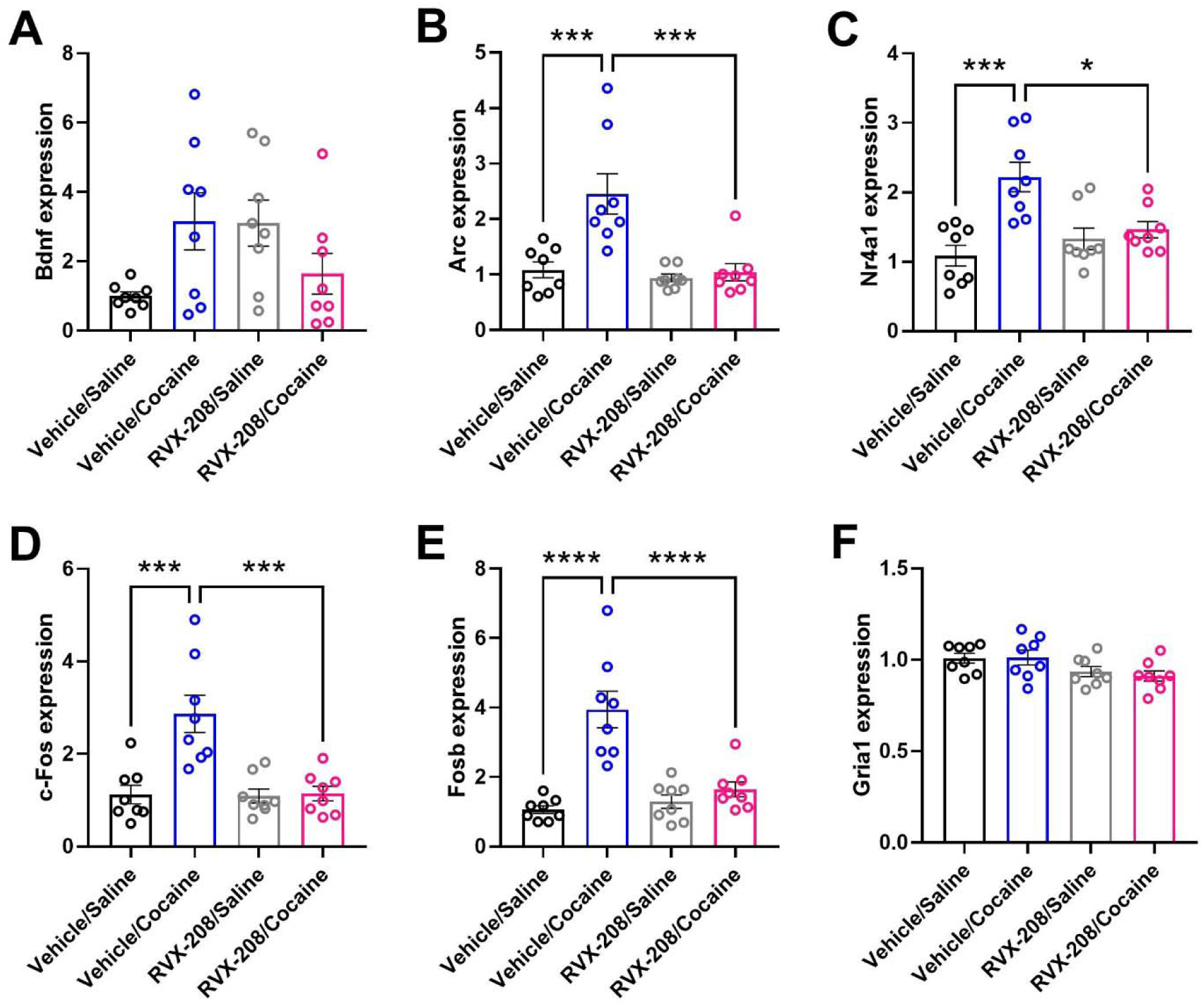
RVX-208 alters cocaine-induced gene expression in the NAc. Mice were treated with vehicle/saline, vehicle/cocaine, RVX-208/saline, or RVX-208/cocaine. Following treatment, (A) *Bdnf*, (B) *Arc* (C), *Nr4a1* (D), *c-Fos*, (E) *Fosb*, and (F) *Gria1* expression levels were measured via RT-qPCR. \**P* < 0.05, \*\*\**P* < 0.001, and \*\*\*\**P* < 0.0001 indicate a significant difference using Bonferroni *post hoc* test. Data are mean ±SEM.

### Effects of RVX-208 on cocaine conditioned place preference and other behaviors

To examine the effects of RVX-208 on cocaine-induced behavior, male mice were injected with vehicle, RVX-208, or JQ1 prior to each cocaine conditioning session. RVX-208 dose-dependently reduced the acquisition of cocaine conditioned place preference (CPP) compared to vehicle treated mice (One-way ANOVA main effect: F _(4, 42)_ = 5.1, P = 0.002 followed by Bonferroni post-hoc test, P = 0.005 for vehicle vs. RVX-208 25 mg/kg and P = 0.03 for vehicle vs. RVX-208 50 mg/kg) (**Figure 5A**). Consistent with previous results, JQ1 (50 mg/kg) attenuated acquisition of cocaine CPP compared to vehicle (Bonferroni post-hoc test, P = 0.002 for vehicle vs. JQ1 50 mg/kg) (**Figure 5A**). In other mice, an acute injection of RVX-208 (50 mg/kg) did not alter distance traveled (t_14_ = 0.35, P = 0.73) (**Figure 5B**) or time spent in the inner and outer zones in an open field compared to vehicle (Two-way ANOVA time in zone effect: F _(1, 7)_ = 151.1, P < 0.0001; treatment effect: F _(1, 7)_ = 2.333, P = 0.1705; interaction effect: F _(1, 7)_ = 0.1335, P = 0.7256) (**Figure 5C**). In the novel object recognition (NOR) task, time exploring the objects during acquisition was not statistically different between the treatment groups (t_13_ = 0.59, P = 0.57) (**Figure 5D**). During the NOR test, vehicle and RVX-208 treated mice showed an increase in time exploring the novel object (NO) compared to the familiar object (FO), but no difference between treatment groups was observed (Two-way ANOVA effect on object exploration time: F _(1, 13)_ = 24.98, P = 0.0002 followed by Bonferroni post-hoc test, P = 0.01 for vehicle FO vs. NO, P = 0.004 for RVX-208 FO vs. NO; treatment effect, F _(1, 13)_ = 0.40, P = 0.5402; interaction effect, F _(1, 13)_ = 0.33, P = 0.5780) (**Figure 5E**). Finally, no significant difference in the novel object discrimination index was observed between the treatment groups (t_13_ = 0.57, P = 0.58) (**Figure 5F**).

**Figure 5.**
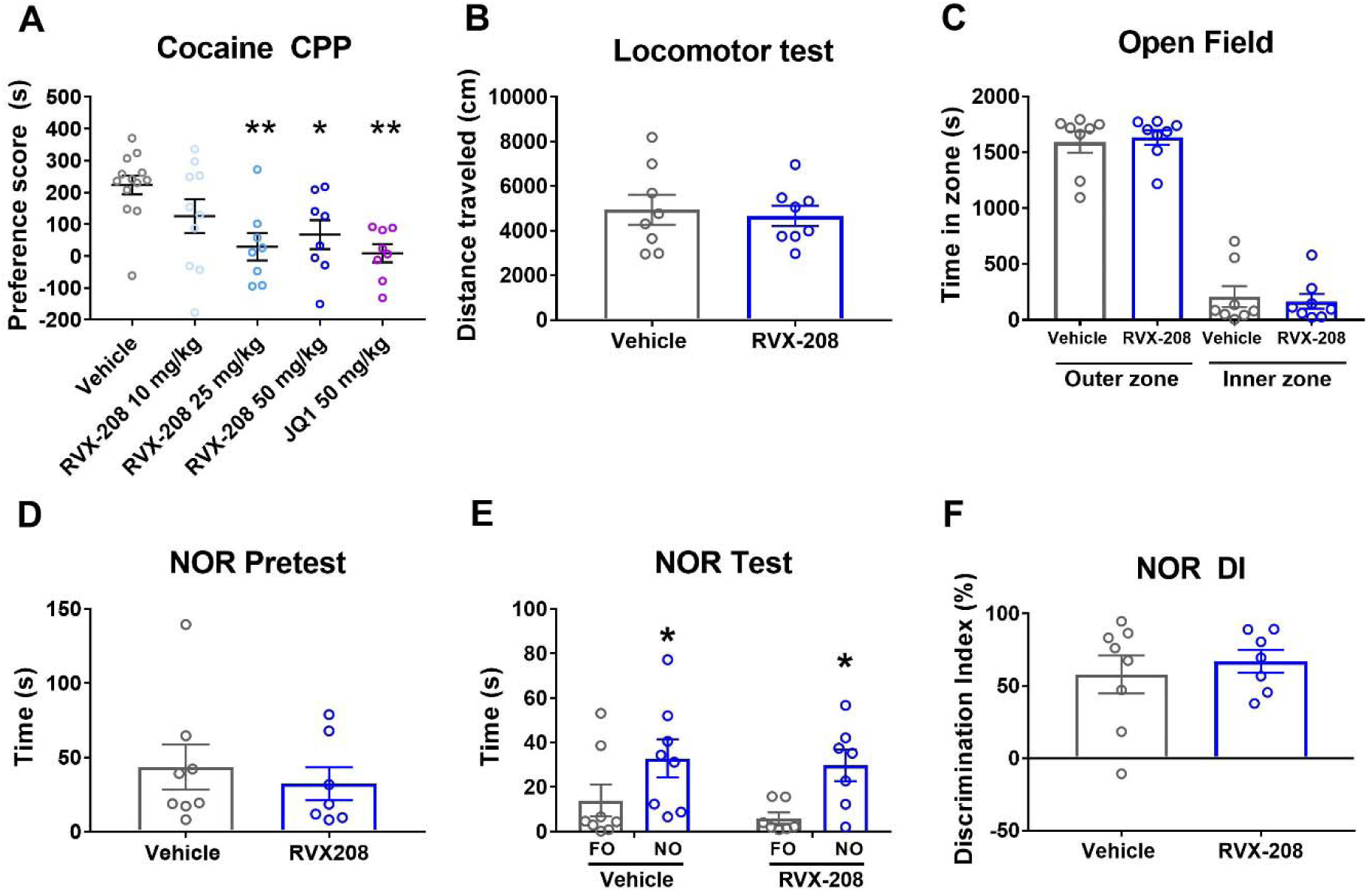
Effects of RVX-208 on behavior. (A) Dose-dependent reduction of cocaine conditioned place preference (CPP) by RVX-208 compared to vehicle-treated mice. (B) The effects of RVX-208 (50 mg/kg) on locomotor activity and (C) time spent in the inner and outer zones in open field. (D) Time spent exploring objects during novel object recognition (NOR) pretest. (E) Time spent exploring the novel object (NO) and familiar object (FO) during the NOR test in vehicle and RVX-208 (50 mg/kg) treated mice. (F) NOR discrimination index (DI) values for vehicle and RVX-208 treated mice. \**P* < 0.05 and \*\**P* < 0.01 indicate a significant difference using Bonferroni *post hoc* test. Data are mean ±SEM.

## Discussion

### A role for BD2-selective BET inhibition in cocaine-induced behavioral and transcriptional responses

In previous studies, we showed that BET protein activity and expression are altered by cocaine use, and BET bromodomain inhibition attenuated behavioral and transcriptional responses to cocaine (Sartor et al., 2015). Consistent with these initial findings, more recent studies have confirmed that BET inhibitors reduce cocaine- and opioid-seeking behaviors (Egervari et al., 2017; Guo et al., 2020; Sartor et al., 2019). To date, however, all addiction-related experiments have used the pan-BET inhibitor JQ1, a tool compound that is not suitable for clinical testing due to its poor pharmacokinetic properties (Tanaka et al., 2016). In addition to having side effects that may restrict their clinical potential as a SUD treatment, pan-BET inhibitors similar to JQ1 block both bromodomains within all BET proteins, restricting our understanding of domain-specific BET mechanisms in drug-seeking behaviors. In the pursuit of more selective and safer therapeutic approaches for SUD, here, we report that RVX-208, a clinically tested, BD2-selective BET inhibitor, attenuated cocaine-conditioned responses without altering distance traveled and anxiety-like behavior in an open field. In previous studies, JQ1 reduced novel object recognition (Korb et al., 2015), an indication that pan-BET inhibitors may have undesirable effects on learning and memory. Distinct from these previous results, we found that RVX-208 did not alter novel object recognition, supporting the concept that BD2-selective BET inhibitors have fewer side effects compared to pan-BET inhibitors.

At the transcriptional level, RVX-208 attenuated cocaine-induced expression of *Arc, Nr4a1, c-fos*, and *Fosb* in the NAc, genes that play key roles in cocaine-induced neuroplasticity (Bannon et al., 2002; Carpenter et al., 2020; Cates et al., 2019; Hope, 1998; Penrod et al., 2020; Salery et al., 2017; Walker et al., 2018; Xu, 2008). Importantly, RVX-208 treatment without cocaine had no significant effect on baseline gene expression in the NAc compared to vehicle-treated mice. These results are in contrast to previous studies that showed pan-BET inhibitors (JQ1 and I-BET858) altered baseline expression of multiple neuroplasticity-associated genes in the NAc and primary neurons (Korb et al., 2015; Sartor et al., 2015; Sullivan et al., 2015). Our current results are in line with previous reports demonstrating that the BET BD2 domain is involved in stimulus-induced gene expression, whereas the BD1 domain (a target of pan-BET inhibitors) maintains steady-state gene expression (Gilan et al., 2020). Similar to our *in vivo* findings, RVX-208 had little effect on gene expression in non-stimulated primary neurons, but it did reduce *Arc* and *Nr4a1* expression in BDNF, but not AMPA, stimulated neurons. JQ1, on the other hand, produced a sustained and robust change in many of the genes measured in non-stimulated and stimulated neurons. When comparing RVX-208 to JQ1, a difference in *c-fos* and *Fosb* expression was also observed. In the current studies, JQ1 increased *c-fos* expression in stimulated and non-stimulated neurons, and in previous studies, JQ1 and I-BET858 elevated *c-fos* and *Fosb* in the NAc and primary neurons (Sartor et al., 2015; Sullivan et al., 2015). In the current studies, RVX-208 reduced *c-fos* and *Fosb* in the NAc of cocaine-treated mice and had no effect on *c-fos* in stimulated and non-stimulated primary neurons. These data indicate that inhibition of the BD1 domain via pan-BET inhibition may be responsible for the observed elevation of *c-fos* and *Fosb*. Future studies are needed to uncover the mechanism of action of this effect. Together, these data reveal that the BET BD2 domain regulates gene expression in stimulated neurons and brain tissue, but unlike pan-BET inhibition, BET BD2 inhibition has minimal impact on baseline expression.

The limited effects on gene expression in non-stimulated cells demonstrates the improved safety profile of BD2-selective inhibitors compared to pan-BET inhibitors. For example, in multiple clinical trials, RVX-208 was shown to be well tolerated in patients without causing thrombocytopenia and gastrointestinal toxicity (Nicholls et al., 2011; Tsujikawa et al., 2019), which are dose-limiting side effects reported in humans treated with pan-BET inhibitors (Amorim et al., 2016; Berthon et al., 2016). In preclinical and clinical studies, RVX-208’s primary indication has been for the treatment of cardiovascular and metabolic diseases (e.g., atherosclerosis, hypertension, hyperlipidemia) by elevating the expression of apolipoprotein AI (ApoA-I) and high-density lipoprotein (HDL) and reducing the expression of inflammation factors (Gilham et al., 2016; Tsujikawa et al., 2019). Interestingly, chronic cocaine use is also known to cause deleterious effects on the cardiovascular system, and cocaine users have an increased risk of developing coronary atherosclerosis (Bachi et al., 2017). Thus, beyond reducing drug-seeking behaviors, RVX-208 may be beneficial to the cardiovascular health of patients suffering from cocaine use disorder.

### Regulation of stimulus-dependent transcription by the BET BD2 domain

Since the development of RVX-208, more selective BD2 BET inhibitors have been reported (Faivre et al., 2020; Gilan et al., 2020). For example, potent and highly selective BD1 (GSK778) and BD2 BET (GSK046) inhibitors were recently identified (Gilan et al., 2020). GSK046 displayed >300-fold selectivity for BD2 over BD1, whereas GSK778 showed ≥130-fold selectivity for BD1 over BD2. The BD1-selective inhibitor GSK778 exhibited similar transcriptional effects compared to pan-BET inhibitors in cancer cells, consistent with previous studies showing that BD1 plays the dominant role in maintaining established transcriptional programs (Picaud et al., 2013). However, distinct from BD1-selective and pan-BET inhibitors, the BD2-selective inhibitor GSK046 primarily affected stimulus-induced gene expression and caused limited disruptions in cell proliferation, cell viability, and BRD4 chromatin binding. In SUD-related research, future studies examining potential differences between modestly selective (RVX-208) versus highly selective (GSK046) BD2 BET inhibitors in cocaine-induced transcriptional and behavioral responses would be of high interest and could potentially guide BET drug development efforts for cocaine use disorder.

Biochemical analysis of the BET protein, BRD4, also indicates that the BD2 domain is involved in stimulus-dependent transcriptional regulation (Chiang, 2016). For example, when the flanking regions of BD2 are dephosphorylated by protein phosphatase 2 (PP2A), the BD2 domain on BRD4 is sterically impeded, preventing it from interacting with acetylated histones and non-histone proteins (Wu et al., 2013). When these regions are phosphorylated by casein kinase 2 (CK2), protein kinase A (PKA), and potentially other kinases, BRD4 undergoes a conformational change that uncovers BD2 and promotes its interactions with other proteins. Though the upstream receptor signaling mechanisms responsible for BRD4 phosphorylation have yet to be fully explored, cocaine and BDNF stimulation were found to increase BRD4 phosphorylation levels in the NAc and primary neurons, respectively (Guo et al., 2020; Korb et al., 2015). The data reported here also suggest that upstream signaling mechanisms following BDNF and cocaine, but not AMPA, stimulation regulate transcription of multiple BET-targeted genes in a BD2-dependent manner.

### Outlook of BD2-selective BET inhibitors for the treatment of cocaine use disorder

Identifying safe and effective treatments for cocaine addiction is an urgent need as there are currently no FDA-approved medications for cocaine use disorder. Developing a brand-new drug treatment, however, takes an enormous amount of time, effort, and money due to years of preclinical and clinical testing for safety and efficacy (Morgan et al., 2011). RVX-208 has been shown to be extremely well tolerated by patients, with safety data now surpassing 2,700 patient-years (Tsujikawa et al., 2019). To date, the only dose-limiting side effect observed with RVX-208 is a reversible and transient elevation in liver enzymes (alanine aminotransferases/aspartate aminotransferases) in a small percentage of patients (Nicholls et al., 2011). With a well-established safety profile in humans, RVX-208 has the potential to make a rapid impact as an epigenetic pharmacotherapy for cocaine use disorder. However, before it can be considered a promising treatment option, RVX-208 must be tested in more sophisticated and translatable models of cocaine use disorder (e.g., short-, long-, and intermittent-access cocaine self-administration and reinstatement/behavioral economic procedures). Additional behavioral tests in female rodents are also needed, as all addiction-related BET studies have been conducted in male rats and mice (Egervari et al., 2017; Guo et al., 2020; Sartor et al., 2015). Finally, while we have identified a few target genes that potentially mediate RVX-208’s effects on cocaine-seeking behavior, future transcriptome-wide studies are necessary to fully elucidate the mechanisms by which RVX-208 regulates cocaine-induced neurobehavioral adaptations. Ongoing efforts to address these questions will determine the therapeutic potential of domain-selective BET inhibitors for the treatment of cocaine use disorder.

## Acknowledgements

This work was supported by National Institute on Drug Abuse grant R00DA040744.

